## Supplementary material for "RCFGL: Rapid Condition adaptive Fused Graphical Lasso and application to modeling brain region co-expression networks": S1 Supplementary

February 8, 2022

#### 1 Theorem for detecting block-diagonal structure

**Theorem 1.** *Denote the set of $p$ genes by $C$. Suppose, $p_1$ many genes belong to a set $C_1$ and $p_2$ many belong to a set $C_2$. And, $C_1 \cap C_2 = \phi$ and $C_1 \cup C_2 = C$. For the genes in $C_1$ to be completely disconnected from those in $C_2$ in each of the resulting network estimates using RCFGL, it is sufficient that $|n_k \mathbf{S}_{ij}^{(k)}| < \lambda_1$ for $k = 1, 2, \cdots, K$ and $i \in C_1, j \in C_2$.*

*Proof.* Suppose there are $K = 2$ classes. By the Karush-Kuhn-Tucker (KKT; [1]) conditions, a necessary and sufficient set of conditions for $\{\Theta\}$ to be the solution to the RCFGL problem is that,

$$\begin{aligned} 0 &= n_1(\boldsymbol{\Theta}^1)^{-1} - n_1\mathbf{S}^{(1)} - \lambda_1\Gamma_1 - \lambda_2\mathbf{W} \odot \Upsilon \\ 0 &= n_2(\boldsymbol{\Theta}^2)^{-1} - n_2\mathbf{S}^{(2)} - \lambda_1\Gamma_2 + \lambda_2\mathbf{W} \odot \Upsilon \end{aligned} \tag{1}$$

where $\Gamma_{1,ij}, \Gamma_{2,ij}$ are the sub-gradients of $|\theta_{ij}^{(1)}|$, $|\theta_{ij}^{(2)}|$ w.r.t $\theta_{ij}^{(1)}, \theta_{ij}^{(2)}$ respectively, and $\Upsilon_{ij}$ is the sub-gradient of $|\theta_{ij}^{(1)} - \theta_{ij}^{(2)}|$ w.r.t $\theta_{ij}^{(1)}$. Consider the matrices,

$$\boldsymbol{\Theta}^{(1)} = \begin{bmatrix} \boldsymbol{\Theta}_1^{(1)} & \mathbf{0} \\ \mathbf{0} & \boldsymbol{\Theta}_2^{(1)} \end{bmatrix}, \boldsymbol{\Theta}^{(2)} = \begin{bmatrix} \boldsymbol{\Theta}_1^{(2)} & \mathbf{0} \\ \mathbf{0} & \boldsymbol{\Theta}_2^{(2)} \end{bmatrix} \tag{2}$$

where $\boldsymbol{\Theta}_1^{(1)}, \boldsymbol{\Theta}_1^{(2)}$ solve the RCFGL problem on the genes in $C_1$, and where $\boldsymbol{\Theta}_2^{(1)}, \boldsymbol{\Theta}_2^{(2)}$ solve the RCFGL problem on the genes in $C_2$. Inspecting Eq. 2, $\boldsymbol{\Theta}^{(1)}, \boldsymbol{\Theta}^{(2)}$ solves the entire RCFGL problem if and only if for all $i \in C_1, j \in C_2$, there exist $\Gamma_{1,ij}, \Gamma_{2,ij}, \Upsilon_{ij} \in [-1, 1]$ such that,

$$\begin{aligned} -n_1\mathbf{S}_{ij}^{(1)} - \lambda_1\Gamma_{1,ij} - \lambda_2 w_{ij}^{12} \cdot \Upsilon_{ij} = 0 \\ -n_2\mathbf{S}_{ij}^{(2)} - \lambda_1\Gamma_{2,ij} + \lambda_2 w_{ij}^{12} \cdot \Upsilon_{ij} = 0 \end{aligned} \tag{3}$$

So, we need to find out the solutions of $\Gamma_{1,ij}, \Gamma_{2,ij}, \Upsilon_{ij}$ that would satisfy Eq. 3. If we set $\Upsilon_{ij} = 0$, Eq. 3 will reduce to,

$$-n_1\mathbf{S}_{ij}^{(1)} - \lambda_1\Gamma_{1,ij} = 0 \quad \text{and} \quad -n_2\mathbf{S}_{ij}^{(2)} - \lambda_1\Gamma_{2,ij} = 0$$

giving the solutions to $\Gamma_{1,ij}, \Gamma_{2,ij}$ as $\Gamma_{1,ij} = -\frac{n_1}{\lambda_1}\mathbf{S}_{ij}^{(1)}$ and $\Gamma_{2,ij} = -\frac{n_2}{\lambda_1}\mathbf{S}_{ij}^{(2)}$. If the inequalities: $|n_1\mathbf{S}_{ij}^{(1)}| < \lambda_1$, $|n_2\mathbf{S}_{ij}^{(2)}| < \lambda_1$ hold, it is obvious that $\Gamma_{1,ij}, \Gamma_{2,ij} \in [-1, 1]$. Thus, $\boldsymbol{\Theta}^{(1)}$, $\boldsymbol{\Theta}^{(2)}$ from Eq. 1 will also be the solution of the entire RCFGL under those two particular inequalities. The proof is similar for $K > 2$ classes and is omitted here. □

### 2 Connection between FGL and FMGL penalties

$$\mathrm{P}^{\mathrm{FGL}}(\mathbf{\Theta}, \lambda_1, \lambda_2) = \lambda_1 \sum_{i \neq j} \sum_{k=1}^{K} |\mathbf{\Theta}_{ij}^{(k)}| + \lambda_2 \sum_{i \neq j} \sum_{k<k'}^{K} |\mathbf{\Theta}_{ij}^{(k)} - \mathbf{\Theta}_{ij}^{(k')}|;$$

$$\mathrm{P}^{\mathrm{FMGL}}(\mathbf{\Theta}, \lambda_1, \lambda_2) = \lambda_1 \sum_{i \neq j} \sum_{k=1}^{K} |\mathbf{\Theta}_{ij}^{(k)}| + \lambda_2 \sum_{i \neq j} \sum_{k=1}^{K-1} |\mathbf{\Theta}_{ij}^{(k)} - \mathbf{\Theta}_{ij}^{(k+1)}|;$$

For $K = 3$, we investigate the second term of $\mathrm{P}^{\mathrm{FGL}}(\mathbf{\Theta})$ focusing on the $ij$-th summand at a time,

$$\begin{aligned}
\sum_{k<k'}^{3} |\mathbf{\Theta}_{ij}^{(k)} - \mathbf{\Theta}_{ij}^{(k')}| &= |\mathbf{\Theta}_{ij}^{(1)} - \mathbf{\Theta}_{ij}^{(2)}| + |\mathbf{\Theta}_{ij}^{(2)} - \mathbf{\Theta}_{ij}^{(3)}| + |\mathbf{\Theta}_{ij}^{(1)} - \mathbf{\Theta}_{ij}^{(3)}| \\
&\leq |\mathbf{\Theta}_{ij}^{(1)} - \mathbf{\Theta}_{ij}^{(2)}| + |\mathbf{\Theta}_{ij}^{(2)} - \mathbf{\Theta}_{ij}^{(3)}| + |\mathbf{\Theta}_{ij}^{(1)} - \mathbf{\Theta}_{ij}^{(2)}| + |\mathbf{\Theta}_{ij}^{(2)} - \mathbf{\Theta}_{ij}^{(3)}| \\
&= 2|\mathbf{\Theta}_{ij}^{(1)} - \mathbf{\Theta}_{ij}^{(2)}| + 2|\mathbf{\Theta}_{ij}^{(2)} - \mathbf{\Theta}_{ij}^{(3)}| \\
&= 2 \sum_{k=1}^{2} |\mathbf{\Theta}_{ij}^{(k)} - \mathbf{\Theta}_{ij}^{(k+1)}|
\end{aligned}$$

We have used the triangle inequality: $|\mathbf{\Theta}_{ij}^{(1)} - \mathbf{\Theta}_{ij}^{(3)}| \leq |\mathbf{\Theta}_{ij}^{(1)} - \mathbf{\Theta}_{ij}^{(2)}| + |\mathbf{\Theta}_{ij}^{(2)} - \mathbf{\Theta}_{ij}^{(3)}|$. Summing all $ij$ terms we get,

$$\sum_{i \neq j} \sum_{k<k'}^{3} |\mathbf{\Theta}_{ij}^{(k)} - \mathbf{\Theta}_{ij}^{(k')}| \leq 2 \sum_{i \neq j} \sum_{k=1}^{2} |\mathbf{\Theta}_{ij}^{(k)} - \mathbf{\Theta}_{ij}^{(k+1)}|.$$

And, finally we establish the connection between $\mathrm{P}^{\mathrm{FGL}}$ and $\mathrm{P}^{\mathrm{FMGL}}$ as,

$$\begin{aligned}
\mathrm{P}^{\mathrm{FGL}}(\mathbf{\Theta}, \lambda_1, \lambda_2) &= \lambda_1 \sum_{i \neq j} \sum_{k=1}^{3} |\mathbf{\Theta}_{ij}^{(k)}| + \lambda_2 \sum_{i \neq j} \sum_{k<k'}^{3} |\mathbf{\Theta}_{ij}^{(k)} - \mathbf{\Theta}_{ij}^{(k')}| \\
&\leq \lambda_1 \sum_{i \neq j} \sum_{k=1}^{3} |\mathbf{\Theta}_{ij}^{(k)}| + 2\lambda_2 \sum_{i \neq j} \sum_{k=1}^{2} |\mathbf{\Theta}_{ij}^{(k)} - \mathbf{\Theta}_{ij}^{(k+1)}| \\
&= \mathrm{P}^{\mathrm{FMGL}}(\mathbf{\Theta}, \lambda_1, 2\lambda_2).
\end{aligned}$$

For $K > 3$, using similar idea we can find a crude bound,

$$\mathrm{P}^{\mathrm{FGL}}(\mathbf{\Theta}, \lambda_1, \lambda_2) \leq \mathrm{P}^{\mathrm{FMGL}}(\mathbf{\Theta}, \lambda_1, \left\lfloor \frac{K^2}{4} \right\rfloor \lambda_2).$$

### 3 Connection between CFGL and RCFGL penalties

$$\mathrm{P}^{\mathrm{CFGL}}(\mathbf{\Theta}, \lambda_1, \lambda_2, \mathbf{W}) = \lambda_1 \sum_{i \neq j} \sum_{k=1}^{K} |\mathbf{\Theta}_{ij}^{(k)}| + \lambda_2 \sum_{i \neq j} \sum_{k<k'}^{K} \mathbf{w}_{ij}^{(kk')} |\mathbf{\Theta}_{ij}^{(k)} - \mathbf{\Theta}_{ij}^{(k')}|;$$

$$\mathrm{P}^{\mathrm{RCFGL}}(\mathbf{\Theta}, \lambda_1, \lambda_2, \mathbf{W}) = \lambda_1 \sum_{i \neq j} \sum_{k=1}^{K} |\mathbf{\Theta}_{ij}^{(k)}| + \lambda_2 \sum_{i \neq j} \sum_{k=1}^{K-1} \mathbf{w}_{ij}^{(kk+1)} |\mathbf{\Theta}_{ij}^{(k)} - \mathbf{\Theta}_{ij}^{(k+1)}|;$$

For $K = 3$, we investigate the second term of $\mathrm{P}^{\mathrm{FGL}}(\mathbf{\Theta})$ focusing on the $ij$-th summand at a time,

$$\begin{aligned}
\sum_{k<k'}^{3} \mathbf{w}_{ij}^{(kk')} |\mathbf{\Theta}_{ij}^{(k)} - \mathbf{\Theta}_{ij}^{(k')}| &= \mathbf{w}_{ij}^{(12)} |\mathbf{\Theta}_{ij}^{(1)} - \mathbf{\Theta}_{ij}^{(2)}| + \mathbf{w}_{ij}^{(23)} |\mathbf{\Theta}_{ij}^{(2)} - \mathbf{\Theta}_{ij}^{(3)}| + \mathbf{w}_{ij}^{(13)} |\mathbf{\Theta}_{ij}^{(1)} - \mathbf{\Theta}_{ij}^{(3)}| \\
&\leq (\mathbf{w}_{ij}^{(12)} + \mathbf{w}_{ij}^{(13)}) |\mathbf{\Theta}_{ij}^{(1)} - \mathbf{\Theta}_{ij}^{(2)}| + (\mathbf{w}_{ij}^{(23)} + \mathbf{w}_{ij}^{(13)}) |\mathbf{\Theta}_{ij}^{(2)} - \mathbf{\Theta}_{ij}^{(3)}| \\
&= \mathbf{w}_{ij}^{*(12)} |\mathbf{\Theta}_{ij}^{(1)} - \mathbf{\Theta}_{ij}^{(2)}| + \mathbf{w}_{ij}^{*(23)} |\mathbf{\Theta}_{ij}^{(2)} - \mathbf{\Theta}_{ij}^{(3)}| \\
&= \sum_{k=1}^{2} \mathbf{w}_{ij}^{*(kk+1)} |\mathbf{\Theta}_{ij}^{(k)} - \mathbf{\Theta}_{ij}^{(k+1)}|.
\end{aligned}$$

where, $\mathbf{w}_{ij}^{*(12)} = (\mathbf{w}_{ij}^{(12)} + \mathbf{w}_{ij}^{(13)})$ and $\mathbf{w}_{ij}^{*(23)} = (\mathbf{w}_{ij}^{(23)} + \mathbf{w}_{ij}^{(13)})$. Define two modified weight matrices as, $\mathbf{W}^{*(12)} = [[\mathbf{w}_{ij}^{*(12)}]]$ and $\mathbf{W}^{*(23)} = [[\mathbf{w}_{ij}^{*(23)}]]$. Denoting $\mathbf{W}^*$ to be the set of the modified weight matrices: $\mathbf{W}^* = \{\mathbf{W}^{*(12)}, \mathbf{W}^{*(23)}\}$, we arrive at the following inequality,

$$\begin{aligned}
\mathrm{P}^{\mathrm{CFGL}}(\mathbf{\Theta}, \lambda_1, \lambda_2, \mathbf{W}) &= \lambda_1 \sum_{i \neq j} \sum_{k=1}^{3} |\mathbf{\Theta}_{ij}^{(k)}| + \lambda_2 \sum_{i \neq j} \sum_{k<k'}^{3} \mathbf{w}_{ij}^{(kk')} |\mathbf{\Theta}_{ij}^{(k)} - \mathbf{\Theta}_{ij}^{(k')}| \\
&\leq \lambda_1 \sum_{i \neq j} \sum_{k=1}^{3} |\mathbf{\Theta}_{ij}^{(k)}| + \lambda_2 \sum_{i \neq j} \sum_{k=1}^{2} \mathbf{w}_{ij}^{*(kk+1)} |\mathbf{\Theta}_{ij}^{(k)} - \mathbf{\Theta}_{ij}^{(k+1)}| \\
&= \mathrm{P}^{\mathrm{RCFGL}}(\mathbf{\Theta}, \lambda_1, \lambda_2, \mathbf{W}^*).
\end{aligned}$$

For $K > 3$, using similar idea we can show that,

$$\mathrm{P}^{\mathrm{FGL}}(\mathbf{\Theta}, \lambda_1, \lambda_2, \mathbf{W}) \leq \mathrm{P}^{\mathrm{RCFGL}}(\mathbf{\Theta}, \lambda_1, \lambda_2, \mathbf{W}^*); \quad \mathbf{W}^{*(kk+1)} = \sum_{r=1}^{k-1} \sum_{n=k+1}^{K} \mathbf{W}^{(rn)} + \sum_{n=k+1}^{K} \mathbf{W}^{(kn)}.$$

where, $\mathbf{W}^*$ is the set of the modified weight matrices, $\mathbf{W}^* = \{\mathbf{W}^{*(kk+1)} : k = 1, 2, \ldots, K-1\}$.

### References

[1] Stephen Boyd, Stephen P Boyd, and Lieven Vandenberghe. *Convex optimization.* Cambridge university press, 2004.

Table 1: **Top pathways detected by the enrichment analysis of the hub-genes found in the common network of the three brain regions.**

| Enrichment FDR | # Hub-genes in Pathway | # Background Genes in Pathway | Fold Enrichment | Pathway |
|---|---|---|---|---|
| 0.00002 | 18 | 40 | 3.69 | Positive regulation of RNA biosynthetic process |
| 0.00002 | 18 | 40 | 3.69 | Positive regulation of nucleic acid-templated transcription |
| 0.00002 | 18 | 40 | 3.69 | Positive regulation of transcription, DNA-templated |
| 0.00002 | 19 | 44 | 3.54 | Positive regulation of nucleobase-containing compound metabolic process |
| 0.00002 | 19 | 42 | 3.71 | Positive regulation of RNA metabolic process |
| 0.00005 | 15 | 30 | 4.10 | Positive regulation of transcription by RNA polymerase II |
| 0.00009 | 18 | 44 | 3.35 | Positive regulation of macromolecule biosynthetic process |
| 0.00019 | 18 | 46 | 3.21 | Positive regulation of cellular biosynthetic process |
| 0.00024 | 18 | 47 | 3.14 | Positive regulation of biosynthetic process |
| 0.00130 | 22 | 73 | 2.47 | Cellular response to chemical stimulus |
| 0.00136 | 7 | 9 | 6.37 | Skeletal muscle tissue development |
| 0.00136 | 7 | 9 | 6.37 | Skeletal muscle organ development |
| 0.00137 | 11 | 22 | 4.10 | Response to organonitrogen compound |
| 0.00158 | 17 | 49 | 2.84 | Regulation of transcription by RNA polymerase II |
| 0.00158 | 19 | 59 | 2.64 | Positive regulation of gene expression |
| 0.00192 | 11 | 23 | 3.92 | Response to nitrogen compound |
| 0.00192 | 17 | 50 | 2.79 | Transcription by RNA polymerase II |
| 0.00415 | 18 | 58 | 2.54 | Cellular response to organic substance |
| 0.00462 | 19 | 64 | 2.43 | Positive regulation of nitrogen compound metabolic process |
| 0.00564 | 6 | 8 | 6.14 | Skeletal muscle cell differentiation |
| 0.00580 | 7 | 11 | 5.21 | Response to mechanical stimulus |
| 0.00761 | 12 | 31 | 3.17 | Response to abiotic stimulus |
| 0.00833 | 13 | 36 | 2.96 | Response to endogenous stimulus |
| 0.00850 | 23 | 91 | 2.07 | Response to chemical |
| 0.00898 | 20 | 74 | 2.21 | Positive regulation of macromolecule metabolic process |
| 0.00898 | 19 | 68 | 2.29 | Positive regulation of cellular metabolic process |
| 0.00898 | 12 | 32 | 3.07 | Cellular response to endogenous stimulus |
| 0.00943 | 7 | 12 | 4.78 | Cellular response to organonitrogen compound |
| 0.00991 | 24 | 99 | 1.99 | Positive regulation of cellular process |
| 0.00992 | 17 | 58 | 2.40 | Tissue development |
